# Molecular resources for the stored grain *Cryptolestes* cryptic pest species (Coleoptera: Laemophloeidae)

**DOI:** 10.64898/2025.12.20.695653

**Authors:** Wee Tek Tay, Melissa C. Piper, Stephen J. Beckett, Daniele Kunz, Paul J. De Barro

**Affiliations:** CSIRO Black Mountain Laboratories, Clunies Ross Street, ACT 2601, Australia; Applied BioSciences, Macquarie University, Sydney, New South Wales, Australia; Department of Biochemistry, University of Cambridge, Tennis Court Road, CB21QW, Cambridge, United Kingdom; CSIRO Ecosciences Precinct, Brisbane QLD, Australia

**Keywords:** Pest interceptions, mitochondrial DNA genomes, molecular diagnostics, flat grain beetles, species genetic gap

## Abstract

Recent research activities relating to evolutionary genetics and molecular characterisation of the stored grain *Cryptolestes* (Ganglbauer) beetle pest species have revealed gaps in public DNA databases, and resulted in molecular diagnostic inconsistencies. This study reports on characterisation of the mitochondrial DNA cytochrome oxidase I (mtCOI) gene from specimens detected during biosecurity inspections at Australia’s border, and surveys of publicly available mtCOI gene sequences, for re-assessment of *Cryptolestes* species status. The results enabled clarification of a new putative *C*. sp. ‘WTT-2016’ species, and supported the close evolutionary relationships between *C*. *ferrugineus* (Stephens)/*C*. *pusilloides* (Steel and Howe), and between *C*. *pusillus* (Schonherr) and the previously identified *C*. sp. ‘WTT-2013’ cryptic species. Widespread confusion between *C*. *ferrugineus*, *C*. *pusiolloides*, *C*. *pusillus* and the two putative novel *C*. sp. ‘WTT-2013’ and *C*. sp. ‘WTT-2016’, highlighted the taxonomic challenge surrounding this beetle genus, while inter-specific genetic distances between *C. spartii* (Curtis)/*C*. *corticinus* (Reitter) suggested they belonged to the same species. Assembled and annotated mitochondrial DNA genomes (mitogenomes) of six *Cryptolestes* species via high throughput sequencing identified assembly errors associated with published mitogenomes of *C. ferrugineus* and *C*. *turcicus* (Grouvelle), and misidentification of *C*. *pusillus*. Based on re-evaluation of intra- and inter-specific genetic distances and congruence from phylogenetic inferences, we proposed a *Cryptolestes* ‘species genetic gap’ at approximately 5% that could help with defining species boundaries. By reconciling these molecular incongruences, this study will contribute to improving agricultural biosecurity preparedness associated with the economically important *Cryptolestes* flat grain beetle species.

## Introduction

Accurate species identification is a major biosecurity consideration and is critical to protecting a nation’s biosecurity in plant health, its pest-free status for global export markets, and agricultural practices. Pests of agricultural importance such as several species of the globally widespread stored grain beetle genus, *Cryptolestes* (the flat grain beetle; FGB), have gained recognition in recent times due to potential resistance to fumigants such as phosphine (Collins, 1998) and difficulties of morphological species identification (Halstead, 1993). This has led to efforts to develop DNA-based species diagnostic methods (Varadinova et al., 2015; Wang et al., 2014), which also included complete mitochondrial DNA genomes (mitogenomes) characterisation (Li et al., 2016; Sun, Li, et al., 2016; Sun, Liu, et al., 2016).

To further complicate species identification, DNA characterisation across a wide range of organisms have detected the presence of species groups separated by significant nucleotide distances but are morphologically indistinguishable (i.e., cryptic species). Cryptic species in pest insect complexes are known (De Barro et al., 2011), where significant genetic differences, life history and behavioural traits including mating incompatibilities existed (Liu et al., 2012). While advances in genomics have significantly contributed to cryptic and putative subspecies identification (Anderson et al., 2016), cryptic species have also been identified by the use of the partial mitochondrial DNA cytochrome oxidase subunit I (mtCOI) gene including in the Coleoptera (Rugman-Jones et al., 2013; Tay et al., 2016).

Detection of insect pests during inspections at the border represents a unique opportunity to identify potential pest origins and arrival pathways, especially when cryptic and/or novel species are involved (e.g., Tay et al., 2022; Handayani et al., 2019). Molecular characterisation of intercepted pest species also offers the unique opportunity of identifying novel and undesirable genetic traits that could potentially be introduced should quarantine fail. Nine *Cryptolestes* species are important with regard to stored grain commodities (Rees, 2004) and are ranked by pest status as major (*C. ferrugineus*), intermediate (*C. turcicus* and *C. pusillus* > *C. pusilloides* and *C. capensis*) and minor (*C. cornutus*, *C. divaricus*, *C. klapperichi*, *C. ugandae*). Of these, Rees (2004) considered *C. ferrugineus* and *C. pusillus* as ‘truly cosmopolitan’, being found in North and South America, Europe and northern Asia, the Mediterranean basin, Africa, South and South East Asia, and Australia and Oceania (see also Berhe et al., 2025). Identification is difficult owing to their small size (i.e., 1.2-2.3mm long), and intra-specific and sexual variation. *Cryptolestes* species identification, therefore, is achieved through examination of genitalia characters (Holloway et al., 2018).

Australia’s grain, pulses, and oilseed industry is forecasted at approximately AUD$26 billion for the 2024/25 season (<u>Grains Australia</u>, accessed 19-December 2025). Much of these commodities end up in bulk storages and so are impacted by pests such as *Cryptolestes*. Although trails using biologicals such as *Beauveria bassiana* entomopathogenic fungus have shown promise for *C. ferruginues* (Tekina-Aydin et al., 2023), control of the flat grain beetle and other stored product pests has relied heavily on phosphine, which is regarded as highly cost effective, and environmentally friendly due to minimal impact on atmospheric ozone. Furthermore, commodities are considered generally safe from chemical residues post fumigation (Chaudhry, 2000). Protecting the effectiveness of phosphine for the grain export industry is, therefore, a high priority as its loss due to insect resistance would in-turn, translate to significant costs for primary producers. The situation might also result in loss of export opportunities under Australia’s policy for ‘nil-tolerance’ of live insects in export shipments (Australian Government, 2024). Incidences of insect resistance to phosphine are known (Konemann et al., 2017), including in *C. ferrugineus* in Australia (Collins, 1998) and in China (Chen et al., 2023). Potential resistance to phosphine/tolerance at a lower level was also suspected in some *C. pusillus* and *C. pusilloides* populations from Queensland’s and New South Wales’s grain producing regions.

While *C. ferrugineus*, *C. pusilloides* and *C. pusillus* are regarded as cosmopolitan species in Australia, a fourth cryptic species (i.e., *C*. sp. ‘WTT-2013’) that is morphologically indistinguishable to *C. pusillus*, but exhibiting limited mtCOI haplotype diversity similar to recent maternal founder events, had been detected in Australia’s eastern regions (Tay et al., 2016). To confirm if *C*. sp. ‘WTT-2013’ likely represents a relatively recent introduced species, molecular characterisation of *Cryptolestes* species being detected through biosecurity inspections at Australia’s border would be a logical starting point. Detection of this cryptic species from intercepted specimens would further indicate its on-going biosecurity risk status. Identifying the associated commodities in which the cryptic species is detected would also improve future pre-border interception efficacies.

Molecular diagnostics of *Cryptolestes* species (Tay et al., 2016; Varadinova et al., 2015; Wang et al., 2014) and characterisation of mitogenomes (i.e., *C. ferrugineus* (Sun, Li, et al., 2016), *C. turcicus* (Sun, Liu, et al., 2016)*, C. pusillus* (Li et al., 2016)), specifically those regularly found in stored grain products, have been reported. The outcomes of these studies clearly demonstrate the significant taxonomic confusion for the *Cryptolestes* genus, and the importance of traditional taxonomy via morphological characters as the foundation to molecular diagnostics (e.g., Polaszek et al., 2025; Conde et al., 2025). Intra-specific mtCOI nucleotide diversity is typically low and varies between 0-2% as reported in diverse species including *Cryptolestes* (<1%, Tay et al., 2016) and other coleopteran species (e.g., Tay et al., 2025a) although at 2.5-3.2% representing potential cryptic species complex has also been reported (Rugman-Jones et al., 2013). This contrasts with studies that report infraspecific nucleotide difference of up to 7-9% for the presumed ‘widely divergent’ lineages of *C. ferrugineus* (Toon et al., 2016) and *C. pusillus* (Pentinsaari et al., 2014; Varadinova et al., 2015), but low as 1-2% at the inter-specific level (*C. ferrugineus* (Sun, Li, et al., 2016), vs. *C. pusillus* (Li et al., 2016)).

In this study, *Cryptolestes* species genetic diversity detected during biosecurity inspections at Australia’s border via the mtCOI molecular diagnostic approach, and comparison between mitogenomes of six *Cryptolestes* species are reported. Potential issues in published *Cryptolestes* mitogenomes are considered and the possibility of significant discordance of DNA-based *Cryptolestes* species identification findings published to-date are discussed to highlight the need to re-evaluate the current *Cryptolestes* species status.

## Materials and Methods

### *Cryptolestes* sample origins

*Cryptolestes* beetle specimens detected during biosecurity inspections at Australia’s border were stored in >90% ethanol and were provided by the Australian Government Department of Agriculture, Fisheries and Forestry (DAFF) in individual specimen vials (Table S1).

### DNA extraction, PCR and Sequencing

Genomic DNA (gDNA) from all *Cryptolestes* individuals intercepted was individually extracted using the Qiagen Blood and Tissue DNA extraction kit and followed the methods of Tay et al. (2016) for PCR and Sanger sequencing using the described primers (CFCOI-F/-R), or the alternative PCR primer pairs Cryp_Barcode_F01 (GTTCATGAGCTGGAATAGCAGGAAC) and Cryp_Barcode_R01 (TAARCCAATYGCTATTATWGCATAA) that amplified 779 base pairs (bp) of the mtCOI molecular diagnostics gene region. The programs Pre-Gap4 and Gap4 (Staden et al., 2000) was used to assemble sequence trace files. All sequences generated were assessed for premature stop codons and insertions/deletions (INDELs) to ensure no nuclear mitochondrial sequences (NuMt/pseudogene) had been included in analyses.

### *Cryptolestes* mitogenomes via HTS, assembly and annotation

The total gDNA from three *Cryptolestes* species (*C. ferrugineus*, n=1; *C. pusillus*, n=1; *C. pusilloides*, n=2) from Australia, one *C. sp. ‘WTT-2013’* (based on partial mtCOI molecular diagnostics; Tay et al. (2016)) found in goods from China, two unknown *Cryptolestes* species (temporarily named *C*. sp. ‘WTT-2016’) found in goods from Vietnam and Malaysia, and one *C. turcicus* found in goods from China (Table S2), were used for constructing species-specific Illumina high throughput sequencing (HTS) genomic libraries following the protocol supplied by the manufacturer. The two randomly selected *C. pusilloides* individuals (T6-01, N4-02) and two randomly selected *C*. sp. ‘WTT-2016’ were chosen to facilitate nucleotide divergence estimates for intra-specific comparison at the mitogenome level. Assembled mitogenomes were annotated using ‘MITOS’ (Bernt et al., 2013) and specifying the invertebrate mtDNA genetic code (5), followed by fine-scale identification of potential start/stop codons for individual protein coding genes (PCGs) within Geneious v11.1.5 (Biomatters Auckland, NZ).

### Comparisons of mtCOI barcoding gene regions of published *Cryptolestes* species

To identify *Cryptolestes* species detected from biosecurity inspections at Australia’s border, all partial mtCOI sequences generated were compared to the GenBank *Cryptolestes* mtCOI sequences. FaBox (Villesen, 2007) was used to identify unique/shared mtCOI haplotypes. The corresponding mtCOI region from complete mitogenomes of *C. ferrugineus* (KT182067; Sun, Li, et al. (2016)), C. *turcicus* (KT070712; Sun, Liu, et al. (2016)) and *C. pusillus* (KT070713; Li et al. (2016)) was not included in the analysis due to issues within these mitogenomes (discussed below). All sequences used were trimmed to 613bp. Gaps at the terminal regions were filled with ‘N’ post trimming, prior to all molecular evolutionary analyses. Nucleotide distance estimates were used in species status delimitation with specific consideration for non-standard nucleotide divergence values across diverse organisms (Cognato, 2006), and that large mtCOI intraspecific nucleotide divergence estimates could be associated with cryptic species (Dinsdale et al., 2010). For the identification of putative *C. ferrugineus*, *C. pusillus*, *C. pusilloides*, *C. turcicus* and the potential cryptic *C*. sp. ‘WTT-2013’, and for the evaluation of *Cryptolestes* species groups from the GenBank DNA database, the representative sequences KF241725, KF241734, KF241739, KF241723, and KF241724 were used.

### Intra- and inter-specific nucleotide diversity estimates

Available GenBank sequences (three complete mtCOI gene sequences and 133 partial mtCOI gene sequences, accessed 07-Jan. 2021) were first identified by Blastn sequence homology search using KF241725 as input query sequence. All GenBank downloaded sequences and those from intercepted specimens were aligned using MAFFT v7.017 (Katoh et al., 2002) default parameter settings (i.e., Scoring matrix: 200PAM/k=2; Gap open penalty: 1.53; and Offset value: 0.123), and trimmed to 613bp (and excluding significantly shorter sequences). Intra- and inter-species nucleotide divergence estimates were calculated using MEGA7 (Kumar et al., 2016) with standard error (s.e.) estimates from 500 bootstrap replications. Nucleotide substitution rates and gamma-shape parameters were also performed within MEGA7, and these parameters were used as input parameters to obtain nucleotide distance estimates using either the Kimura 2-parameter (K2P, (Kimura, 1980)) nucleotide substitution model, or the uncorrected pairwise nucleotide distances (*p*-dist) approach. The s.e. estimates representing the variables were used to obtain the respective 95% confidence intervals (C.I.; calculated as 1.96 x s.e.) and for plotting on the y-axis against the estimated mean genetic distances (i.e., non-variables, x-axis) for visualisation of differences within and between species groups, and to allow the potential genetic gap to be identified to support species delimitation (Dinsdale et al., 2010).

### Mitogenome protein coding gene (PCG) nucleotide divergences in six *Cryptolestes* species

To estimate nucleotide divergences between economically significant FGB species, the putative cryptic *C*. sp. ‘WTT-2013’ and novel *C*. sp. ‘WTT-2016’, all eight mitogenomes were aligned with an outgroup Cucujidae species (MK614530) using MAFFT v7.017. All 13 PCG sequences were extracted for re-alignment and end-trimming. A 71bp region within the NAD6 gene in all individuals was removed due to the missing region in *C.* sp. ‘WTT-2013’ (i.e., NAD6 nt8,715-8,785; Tay et al. (2024)). Aligned concatenated PCG sequences (11,024bp) were used to calculate the *p*-dist prior to construction of a mitogenome concatenated PCG phylogeny. We also calculated the *p*-dist for the complete mtCOI gene for comparison with the concatenated PCGs *p*-dist.

### Phylogenetic analysis

#### A. Partial mtCOI phylogeny

For the inference of phylogenetic relationships, a partial mtCOI maximum likelihood (ML) phylogeny on the N-terminal (i.e., ‘DNA barcode’) region used in the Intra- and inter-specific nucleotide diversity analyses was constructed using IQ-tree (Nguyen et al., 2015). The ‘automatic evolutionary rates’ option via ModelFinder was selected to estimate optimal nucleotide substitution rates and evolutionary model, and specifying 1,000 ultra-fast bootstrap replications to estimate branch node support. Published Cucujidae beetle species’ mtCOI sequences used as out groups included: (i) *Cucujus clavipes* GU176341, (ii) *C*. *haematodes* KM439405, KM441995, (iii) *C*. *cinnaberinus* KM447566, (iv) *Pediacus depressus* KM441566, (v) *Pediacus fuscus* KJ964320, and two unpublished GenBank public records (vi) *Cucujus clavipes* KM847989, and (vii) *Pediacus fuscus* KJ203564. Visualisation of the phylogenetic tree was via FigTree v1.4.0.

#### B. Concatenated mitochondrial protein coding genes phylogeny

The ML phylogenetic relationships of the detected *C*. *ferrugineus*, *C*. *pusilloides*, *C*. *turcicus*, *C*. *pusillus*, *C*. sp. ‘WTT-2013’, and *C*. sp. ‘WTT-2016’ were inferred using 11,024bp of trimmed and aligned concatenated sequences from the 13 mitochondrial DNA PCGs via IQ-tree as detailed, and included the PCGs from the Phalacridae gen. sp. Cucujidae beetle (MK614530; Jin et al. (2020)) as the outgroup. The PCG partition was specified and used as input file to enable optimal evolutionary models and substitution rates to be determined for individual PCGs within IQ-tree. Visualisation of the phylogeny was by Dendroscope 3.

### Analysis of published *Cryptolestes* mitogenomes

Published mitogenomes of *C. ferrugineus* (KT182067), *C. turcicus* (KT070712), and *C. pusillus* (KT070713) were compared against assembled mitogenomes from Tay et al. (2024) through MAFFT alignment to identify potential contig assembly errors and to confirm species status. For potential problematic gene regions in the published mitogenomes, uncorrected *p*-dist estimates were provided against assembled mitogenomes from this study to identify likely contamination sources.

## Results

### *Cryptolestes* species from biosecurity inspections

Between September 2013 and January 2015, a total of 92 interceptions of *Cryptolestes* beetles from contaminated plant commodities (e.g., cashews, peanuts, sorghum, green coffee beans), toys, building materials (e.g., Reed fencing) and plant parts (e.g., rhizomes) were detected during biosecurity inspections at Australia’s international ports. The numbers of beetle adults and larvae intercepted in each item ranged from one to 46 individuals, and involved diverse geographic origins (Table S1, Table S2, Table S3).

Of the six *Cryptolestes* species detected and identified by molecular diagnostics, four (*C*. *ferrugineus*, *C*. *pusilloides*, *C*. *pusillus, C*. sp. ‘WTT-2013’; Tay et al. (2016)) were known to be present in Australia, and two were not known to be present in Australia (i.e., *C*. *turcicus*, and an unknown *C*. sp. ‘WTT-2016’). The *C*. sp. ‘WTT-2016’ specimens were detected in goods that originated from China, Vietnam, Malaysia and Colombia, and was also the most commonly detected species identified by molecular diagnostics. The *C*. sp. ‘WTT-2016’ has been identified as *C. pusilloides* (Hendrich et al., 2015; Wang et al., 2014), but differed significantly from the partial mtCOI sequence of a specimen morphologically identified as *C. pusilloides* (Tay et al., 2016).

### Molecular characterisation of mitogenomes of six *Cryptolestes* species

Eight mitogenomes that belonged to six *Cryptolestes* species (1x *C. ferrugineus*; 2x *C. pusilloides*; 1x *C. pusillus;* 1x *C. turcicus*, 2x *C*. sp. ‘WTT-2016’; 1x *C*. sp. ‘WTT-2013’) were assembled and annotated. The six *Cryptolestes* beetle species’ draft mitogenomes ranged from approximately 15,186bp to 15,341bp, were high in A-T nucleotide composition (Table S2), and had 13 PCGs, 22 tRNA’s, and two rRNA’s genes, similar to mitogenomes of other insects including the Coleoptera. The mitogenome gene orientations for all six *Cryptolestes* species were identical to other reported Cucujidae species.

### Detection of the ‘unknown’ cryptic *C.* sp. WTT-2013

Four of the seven *Cryptolestes* beetles intercepted from rhizomes (PBCRC-08, PBCRC-09; Table S1, Tay et al. (2024)) were successfully identified by molecular diagnostics, having shared high sequence homologies (99%) to the two mtCOI partial gene sequences (KF241724, KJ502178) of the putative *C*. sp. ‘WTT-2013’ detected in Queensland and New South Wales (Tay et al., 2016). Furthermore, there existed 11 other partial/complete mtCOI sequences in GenBank that also showed high sequence homologies to KF241724, KJ502178 (Tay et al., 2016)) and to PBCRC-08, PBCRC-09 (Tay et al., 2024)). Of the total 15 sequences, all but two (KF241724, KJ502178) were from China, and have been assigned as *C*. sp. ‘WTT-2013’ (KF241724, KJ502178), *C. ferrugineus* (KT182067), *C. turcicus* (KT070712), or *C. pusillus* (KT070713, JQ708206, KC977922) (Table S2).

### Detection of the unknown putative *C.* sp. ‘WTT-2016’

A total of 11 intercepted *Cryptolestes* specimens on goods from Malaysia, Colombia, Vietnam, and China (Table S1) shared high sequence homologies but were sufficiently divergent from other *Cryptolestes* species and were all assigned with a putative species status (i.e., *C*. sp. ‘WTT-2016’). The challenge of ascertaining the true identity of *C*. sp. ‘WTT-2016’ is further compounded by sequences from China (KC436315, KC977916) and Germany (KM450594) being reported as *C. pusilloides* (Fig. S1). A survey of literature and from this study showed that *C*. sp. ‘WTT-2016’ likely represents a widespread species present in South America, Asia and Europe.

### *Cryptolestes* species status assessment of GenBank sequences

Assessment the mtCOI N-terminal (i.e., ‘DNA barcode’) region sequences representing the 10 *Cryptolestes* species and including all unique mtCOI haplotypes from this study resulted in 129 sequences being analysed. There were 29 potentially problematic and two unclassified sequences being assigned either as *C. ferrugineus* (n=15), *C. pusillus* (n=9), *C. pusilloides* (n=3), or *C. turcicus* (n=2) (Fig. S1). Furthermore, seven sequences were associated with either *C. corticinus* or *C. spartii*. These two species exhibited low nucleotide distances (average: 3.5% *p*-dist, range 0 to 4.57%; Table S4), indicating potential confusion over their species status. Phylogenetic analyses (discussed below) were used to further support species status assessment. Taken as a whole, approximately 30% (i.e., 39/129) of the publicly available *Cryptolestes* partial mtCOI sequences were associated with taxonomic confusion (Fig. S1).

### Phylogenetic analysis

Based on partial (613bp) mtCOI sequences, all *Cryptolestes* species groups including the putative *C*. sp. ‘WTT-2013’ and *C*. sp. ‘WTT-2016’ were assigned to 10 phylogenetic clades (clades A to J) representing individual species groups with high branch node confidence (80% to 100%; Fig. 1).

**Fig. 1:**
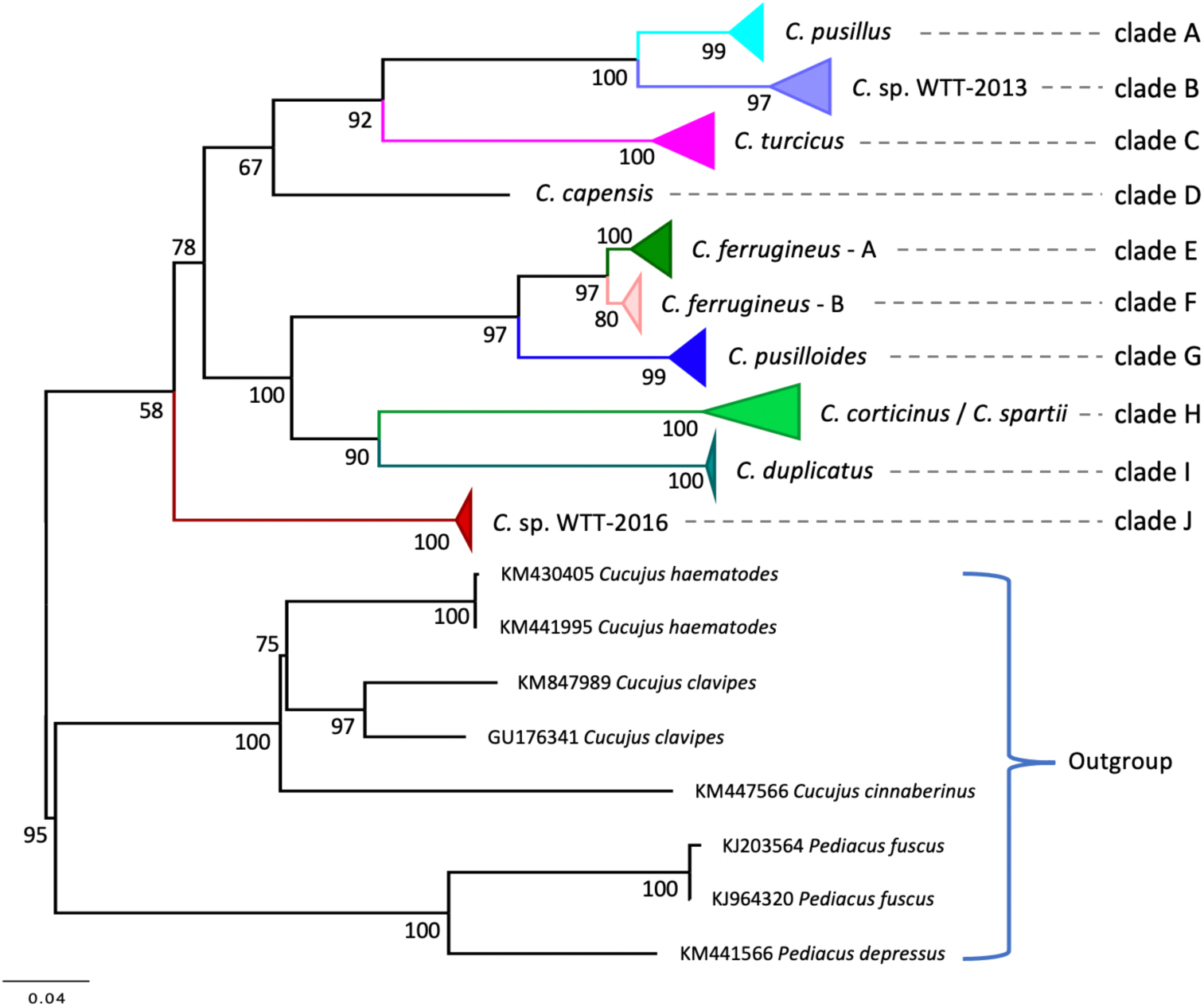
*Cryptolestes* phylogeny based on mtCOI partial gene sequences from GenBank and this study. Refer to Fig. S1 for GenBank accession numbers and to Tay et al. (2024) for sequences used. Note: Best-Fit substitution model (determined using ModelFinder): TIM2+F+I+G4; ML consensus tree log-likelihood score: -5403.4655; Bayesian information criterion score (BIC): 12394.8391; Akaike information criterion score (AIC): 11338.8499.

The major species (i.e., *C. ferrugineus*, *C. pusilloides*, *C. pusillus*, *C. capensis*, *C. turcicus*; Rees (2004)) are paraphyletic to each other. The *C*. sp. ‘WTT-2016’ was basal to other *Cryptolestes* species, and importantly, retained its monophyletic position with *C. pusillus* (Tay et al., 2016). Low bootstrap estimates for the phylogenetic placements of individual clades with respect to each other do not impact on the proposed species status, but rather, reflect the need for more comprehensive species diversity surveys and for greater nucleotide data to better resolve evolutionary relationships between clades. The 10 clades identified are detailed below.

#### Clade A: Cryptolestes pusillus

All eight *C*. *pusillus* haplotypes (i.e., KF241734 -KF241738, KJ502175 -KJ502177; Tay et al. (2016)) and four *C. pusillus* haplotypes from GenBank (KC977917, KC977930, JQ708205, KM449111) clustered with high bootstrap support (91%). Four haplotypes (PBCRC-10, -23, -24, -25) from this study also clustered as *C*. *pusillus*. However, nine GenBank sequences identified as *C*. *pusillus* were not clustered in this clade. These included a complete mitogenome (KT070713), and eight partial mtCOI sequences (KC977918 -KC977923, JQ708206, KC977931) that shared low (91-92%) sequence identity with KF241734 from Tay et al. (2016).

#### Clade B: *Cryptolestes* sp. ‘WTT-2013’

*Cryptolestes* species’ identity in this clade represents one of the most confusing in the public DNA databases. Previously, Tay et al. (2016) argued that, based on pairwise DNA sequence divergence, the haplotypes identified (KJ502178, KF241724) from *C*. sp. ‘WTT-2013’ was likely a cryptic species of *C*. *pusillus*. A study of *Cryptolestes* species diversity from China, USA and the Czech Republic by Wang et al. (2014) also identified highly divergent sequences within *C. pusillus*. However, the authors concluded that intra-specific nucleotide diversity in *C. ferrugineus* and *C. pusillus* varied between 0 and 8.9%, but was low in *C. pusilloides* and *C. turcicus* (max. < 1%).

#### Clade C: Cryptolestes turcicus

Of the published (KF241723, KC977933, JQ708207, KC977913, KT070712) and unpublished (PBCRC-18, -19; China samples from within reed fencing; Tay et al. (2024)) *C*. *turcicus* partial mtCOI sequences analysed, only the mtCOI gene from the full mitogenome (Sun, Liu, et al., 2016) did not cluster in Clade C, and was possibly either due to contig assembly errors or PCR contamination (discussed later).

#### Clade D: Cryptolestes capensis

Two partial mtCOI sequences of *C*. *capensis* (Varadinova et al., 2015; Wang et al., 2014) were available from GenBank. Reproductive organs were used for species identification, however prepared slides of the genitalia characters were not kept (Z. Li, pers. comm.) and therefore could not be independently verified. The species identity of *C. capensis* is tentatively assumed to be accurate.

#### Clades E &F: Cryptolestes ferrugineus

Two *C. ferrugineus* sister clades (E and F) were identified and shared high sequence homologies (average *p*-dist: 2.8%; range: 1.47-3.59%; Table S4). The maximum *p*-dist between clades E and F was less than between closely related species (e.g., *C. pusillus*/*C*. sp. WTT-2013; *C. ferrugineus*/*C. pusilloides*; Fig.1, Fig. S1). Sequences from the *C. ferrugineus* sister clades, (i.e., *C. ferrugineus*-A (17 sequences), *C. ferrugineus*-B (7 sequences)) also clustered together with a high (97%) node confidence value, but excluded 16 ‘*C. ferrugineus*’ published and unpublished sequences that shared lower (91%-92%) sequence homologies to the *C. ferrugineus* Cfer-01 haplotype (KF241725) and likely represented misidentification. For example, Pentinsaari et al. (2014) reported sequences of *C*. *ferrugineus* based on voucher samples (KJ964655, KJ961815) that either clustered confidently with *C*. *ferrugineus* sequences within the *C. ferrugineus* clade (i.e., KJ964655) or with Clade G (*C. pusilloides*; KJ961815). Similar confusion surrounded other voucher *C. ferrugineus* specimens (i.e., KM441850, KM447067; Hendrich et al. (2015)) that also clustered confidently in Clade G (see also Fig. S1).

#### Clade G: Cryptolestes pusilloides

In Clade G, 12 sequences reported in Tay et al. (2016) and from this work (PBCRC-07, -13, -14, -27, Cpld-N4-02, Cpld-T6-01; Tay et al. (2024)), and sequences identified as *C*. *ferrugineus* (JQ708204, KC977927, KC977924, KM441850, KM447067, KJ961815, KX528594, KX528597, KX528588, KX528595, and MG458967 (GenBank Unpublished)) were confidently clustered together (Fig. 1; Fig. S1). The sequences used by Hendrich et al. (2015) (KM450594) and Pentinsaari et al. (2014) (KJ961815) were voucher specimens, further highlighting taxonomic challenges in *Cryptolestes* species including for voucher specimens.

#### Clade H: Cryptolestes spartii/C. corticinus

Two partial mtCOI sequences for *C*. *spartii* (KM450873, KM452565) and five partial mtCOI sequences for *C*. *corticinus* (KM447571, KJ963948, KU918377, KU912013, KJ963948) existed in GenBank. All specimens belonged to museum voucher samples (Hendrich et al., 2015; Pentinsaari et al., 2014). Due to the lack of additional sequences for intra-species comparison for each of these two species, it was not possible to ascertain their accuracy and validity. The *P*-dist estimates between these two species are below that observed for known inter-specific nucleotide distances (Table S4), highlighting the need of species status re-assessment.

#### Clade I: Cryptolestes duplicatus

There were eight sequences from museum voucher *C. duplicatus* specimens (Germany: KM451562, KU919513, KU909614, KU908839, KU912488, KU917587; Belgium: KM440172; Sweden: KJ966304) (Hendrich et al., 2015; Pentinsaari et al., 2014). All *C. duplicatus* sequences were clustered confidently as a clade.

#### Clade J: *Cryptolestes* sp. ‘WTT-2016’ (putative)

The species name of this group of *Cryptolestes* beetles is not known. Of the 14 sequences used in this study, 11 were from beetles collected during biosecurity inspections, and involved commodities from diverse origins. Three other partial mtCOI sequences were identified as *C. pusilloides* (from China: KC977916, KC436315; from Germany: KM450594). *Cryptolestes* sp. ‘WTT-2016’ has therefore been recorded directly, or in goods originating, from Asia (China, Vietnam, Malaysia), Europe (Germany), and South America (Colombia), and represents a species that could successfully establish in Australia due to regular interceptions and its presence in diverse eco-climatic zones.

### Mitogenome phylogeny of *Cryptolestes* species

The phylogeny of the six *Cryptolestes* species based on the mitochondrial PCGs (Fig. 2) was in overall agreement with the partial mtCOI gene phylogeny (i.e., *C. pusilloides*/*C. ferrugineus*, and *C. pusillus*/*C*. sp. ‘WTT-2013’ as closely related sister species). The *C. turcicus* species was basal to the clade that included both *C. pusillus* and *C*. sp. ‘WTT-2013’, while *C*. sp. ‘WTT-2016’ was basal to the *C. turcicus*/*C. pusillus*/*C*. sp. ‘WTT-2013’ clade, and branch node confidence values were also high (90-100%). A major difference between the mitogenome and the partial mtCOI phylogenies is that based on the concatenated PCG sequences, the *C. ferrugineus*/*C. pusilloides* clade was shown to be the most ancestral species clade, whereas in the partial mtCOI phylogeny which included more species, the novel *C*. sp. ‘WTT-2016’ had the most ancestral placement.

**Fig: 2:**
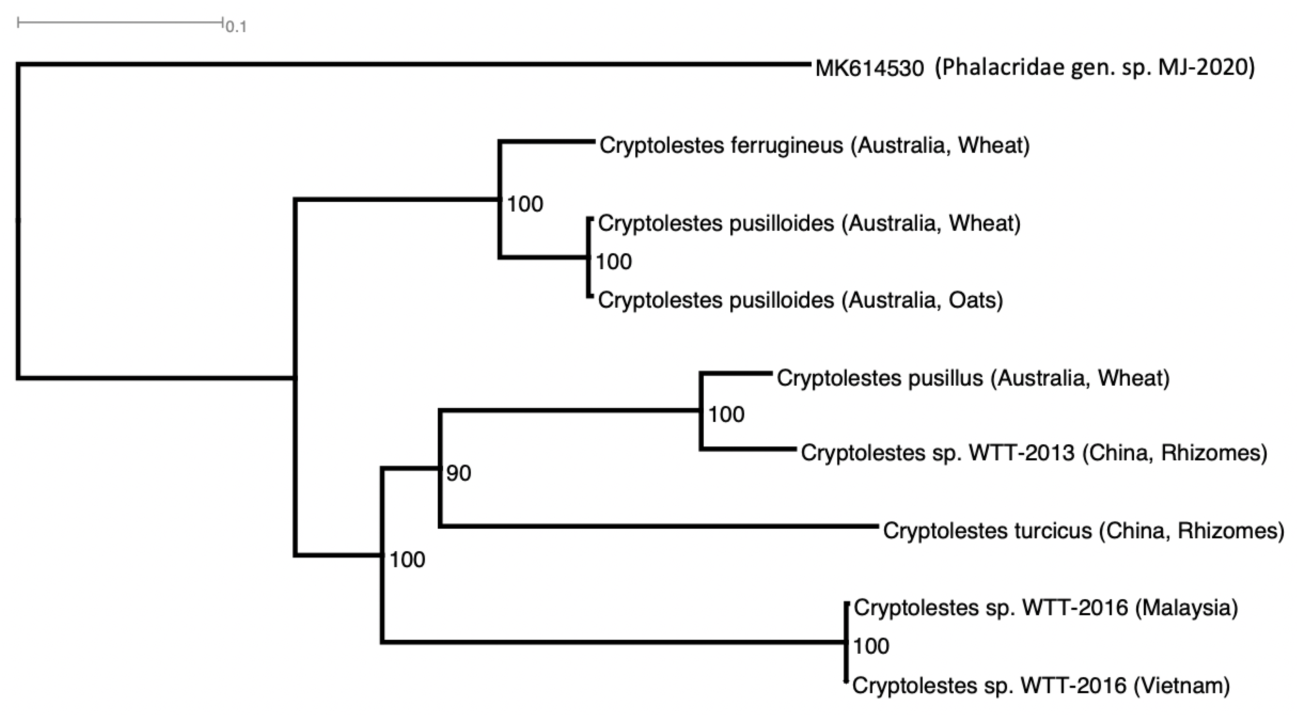
Phylogeny of six *Cryptolestes* species as inferred from concatenation of the 13 mitochondrial protein coding genes, with the published Cucujidae mitogenome MK614530 (Phalacridae gen. sp. MJ-2020) as an outgroup. Note: Best-Fit substitution model (determined using ModelFinder): HKY+F+G4; ML consensus tree log-likelihood score: -41276.4928 (s.e. 315.9798); BIC: 83632.6985; AIC: 82784.9902.

### Nucleotide distances

Intra-specific nucleotide distances estimated for each of the nine *Cryptolestes* species in this study were typically low within each species (e.g., *C. duplicatus*: ranged 0.0 to 0.0049: *C. corticinus*/*C. spartii*: 0.0 to 0.0457; Table S4). Plotting the average between species nucleotide distance estimates based on either the *p*-dist or the K2P evolutionary models showed the K2P estimates to be typically higher than the *p*-dist estimates, although at the intra-specific level both estimates were similar (Fig. 3). A clear nucleotide distance ‘gap’ was evident between the intra- and inter-specific comparisons at approximately 5-6% (range ∼4.5-7.5%). The two *C. ferrugineus* sister clades (i.e., clades E and F; Fig. 1) were assigned as a single species based on partial mtCOI nucleotide distance estimates. Inter-specific nucleotide distances of *C. ferrugineus*/*C. pusilloides* and *C. pusillus*/*C*. sp. ‘WTT-2013’ were approximately 7.67% based on the *p*-dist approach (10.8-12.4% by the K2P model), and further supported them as closely related species (Fig. 3; Table S4). Inter-specific estimates of nucleotide distance between the other *Cryptolestes* species were otherwise large (*p*-dist: 13.78-17.94%; K2P: 15.2-19.4) (Table S4).

**Fig. 3:**
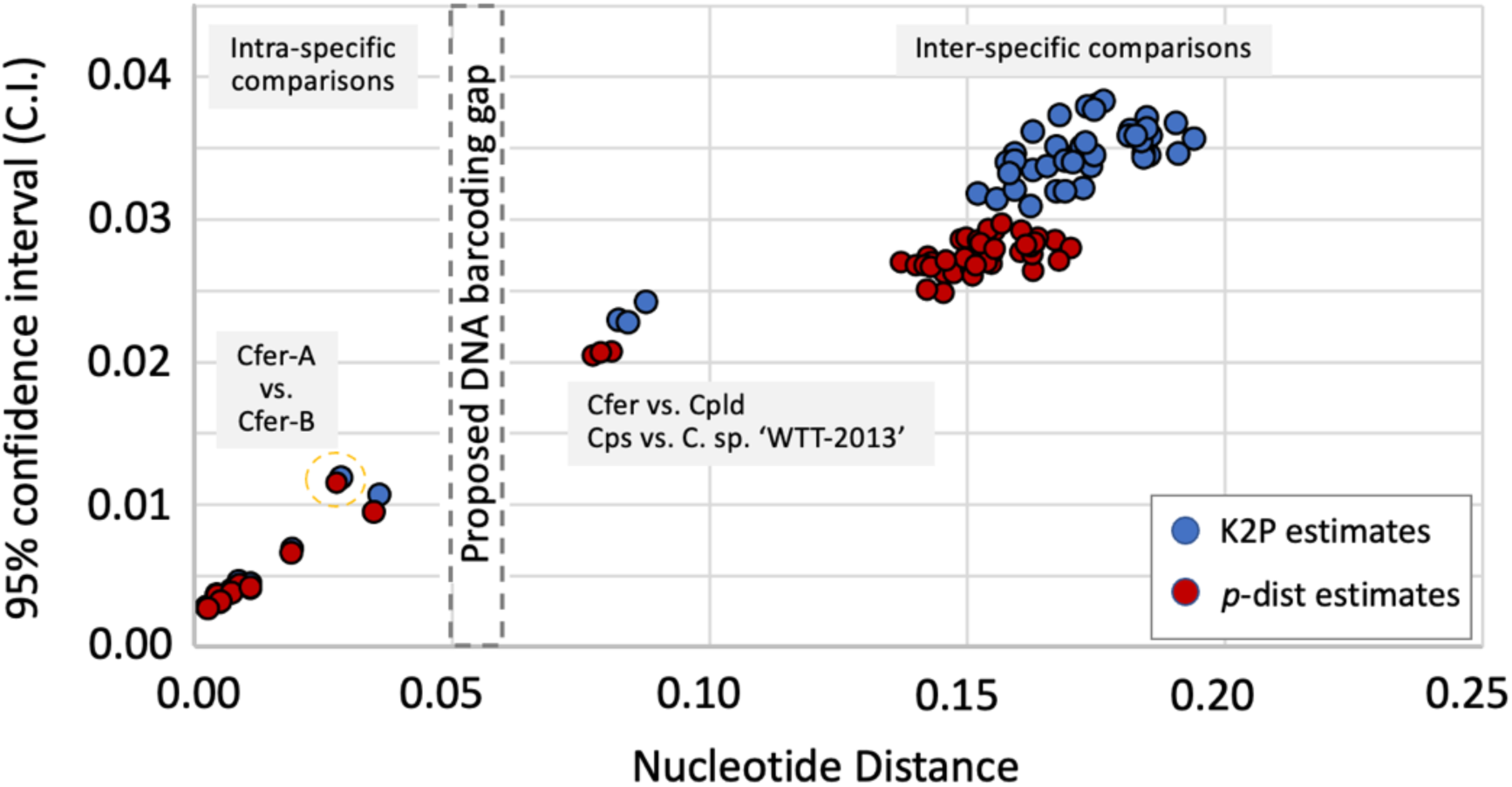
Scatter plots of intra- and inter-specific nucleotide distance estimates of *Cryptolestes* species, showing both expected low and high nucleotide distances for within and between species, respectively. A proposed ‘DNA barcoding gap’ (dashed line box at 5-6%) is shown. Close genetic relationship between *C. ferrugineus*/*C. pusilloides* (Cfer vs. Cpld) and *C. pusillus*/*C*. sp. ‘WTT-2013’ (Cpls vs. *C*. sp. ‘WTT-2013’) is also shown. *C. ferrugineus* sister clades E and F (Fig. 1) are represented by Cfer-A and Cfer-B, respectively (dashed-line circle). Y-axis is the 95% confidence interval (C.I). Average intra-specific nucleotide estimates also includes that between *C. spartii* and *C. corticinus*. Estimates provided were for K2P and *p*-dist models.

### *Cryptolestes* mitogenome PCGs nucleotide distances

Intra-specific nucleotide distances based on concatenated mitogenome PCG sequences (11,024bp) similarly detected low estimates (*p*-dist < 1%) for the two *C*. sp. ‘WTT-2016’ and the two *C. pusilloides*. Inter-specific *p*-dist estimates between closely related species were approximately 6.3-6.8%, while between distantly related *Cryptolestes* species, *p*-dist estimates were generally higher (15-18%, Table S5).

### Analyses of published *Cryptolestes* mitogenomes

Sequence alignment of *C. ferrugineus* (KT182067), *C. turcicus* (KT070712), and *C. pusillus* (KT070713) mitogenomes showed that KT182067 and KT070713 were overall similar (*p*-dist: 2.15%; expected at intra-specific level, Table S5) with the difference largely caused by an approximately 2,300bp nucleotide dissimilarity that spanned the COIII-trnG-ND3-trnA-trnR-trnN-trnS1-trnE-trnF-ND5 gene region. Multiple sequence alignment of KT182067 (*C. ferrugineus*) with assembled mitogenomes of the six species from this study suggested contig assembly errors for this KT182067 mitogenome (Fig. S2 a & b).

Similarly, comparison of the *C. turcicus* mitogenome (KT070712) with mitogenomes from this study indicated KT070712 to be predominantly *C. turcicus*, with the exception of an approximately 700bp region at the mtCOI barcoding gene region, where it matched with high sequence homology to the *C*. sp. ‘WTT-2013’ mtCOI gene. This provided evidence that the published *C. turcicus* mitogenome (KT070712) was a chimera likely caused by contamination/contig assembly errors (Fig. S3 a & b).

Excluding two missing mitogenome regions (i.e., a 93bp region from *C*. sp. ‘WTT-2013’ ND6 gene, and an approximately 407bp mitogenome region from the AT-rich region; Tay et al. (2024)), the published *C. pusillus* mitogenome (KT070713) and the *C*. sp. ‘WTT-2013’ mitogenome (Tay et al., 2024) shared 99.23% sequence identity, suggesting that they were the same species. The species identity of KT070713 and *C*. sp. ‘WTT-2013’ will require confirmation. Based on Tay et al. (2016) however, it would suggest that KT070713 (but excluding the approximately 700bp 5’ end of the mtCOI gene) and *C*. sp. ‘WTT-2013’ are the same cryptic species that is genetically distinct to *C. pusillus*.

## Discussion

In this study, molecular diagnoses of *Cryptolestes* beetles detected during biosecurity inspections at Australia’s border, and re-analyses of publicly available partial mtCOI sequences and mitogenomes, demonstrated the pervasiveness of species misidentification in this agriculturally important beetle genus. The previously detected cryptic *C*. sp. ‘WTT-2013’ in eastern Australia (Tay et al., 2016) is also likely an introduced exotic species with a probable Asian origin. Furthermore, while *C. ferrugineus* and *C. pusillus* have been regarded as the most widespread and agriculturally important species (Rees, 2004), the novel *C*. sp. ‘WTT-2016’ represented the most frequently intercepted species in Australia from diverse agricultural commodities, building material, and personal effects, and potentially has a wide geographic presence in Asia, Europe, and South America, and unknown presence in North America and Africa, as no *Cryptolestes*-contaminated goods were intercepted from these two continents during the sampling period. Given the species was identified from continents with similar latitudes to the North American and African continents, international movements of contaminated goods to or from globally diverse regions could lead to accidental introductions of this species, as exemplified by the recent spread of various highly invasive insect pest species that were shown to be associated with anthropogenic-related activities (e.g., Rugman-Jones et al., 2013; Lopes-Da-Silva et al., 2014; Tay et al., 2016; Elfekih et al., 2018; Rane et al., 2023; Hoffmann et al., 2024; Magalhaes et al., 2025; Tay et al., 2025b).

The confusion in *Cryptolestes* species identification (e.g., Hendrich et al., 2015; Toon et al., 2016; Varadinova et al., 2015; Wang et al., 2014) demonstrate the significant taxonomic challenge in this beetle genus, which also extends to museum voucher specimens. Possible factors underpinning poor species identification may include: (i) the unanticipated presence of cryptic species, (ii) the expectation that voucher *Cryptolestes* species from different museums were in agreement, and (iii) possible laboratory/analytical-related mistakes including potential PCR contaminations or contig assembly errors. Contig assembly errors and failure to cross-check assembled genomes/DNA contigs can exacerbate confusion for molecular diagnostic efforts, with potential significant biosecurity and pest management implications if uncorrected. This is especially important in cryptic and highly invasive insect pest species, and could be factors that underpinned the persistent molecular diagnostics-related misidentification of *Cryptolestes* (Fig. S1).

Various coleopteran partial mtCOI gene studies showed that in beetles, interspecific nucleotide distances ranged from approximately 4.4-12.8% (Aykut et al., 2019; Monaghan et al., 2005). Similar *p*-dist estimates with other gene regions (Machado et al., 2017) and evidence presented here therefore support a need for taxonomic revision for *Cryptolestes* species. This study to characterise intra- and inter-specific nucleotide distances showed that at the within species level, partial mtCOI gene *p*-dist estimates in *Cryptolestes* typically ranged between 0-4.6% for within species, and from approximately 7.5-17.9% for between species (Fig. 3; Table S4), thereby allowing a ‘barcoding gap’ to be tentatively proposed at between approximately 5-6% mtCOI nucleotide distance (Fig. 3), and lending support that *C*. sp. ‘WTT-2013’ is likely to be a cryptic species of *C. pusillus* (Tay et al., 2016) (Fig. S1, Table S4). Phylogenetic analysis based on concatenated mitogenome PCGs also supported the species status of *C*. sp. ‘WTT-2013’, and contrasted with the significant branch length difference observed for intraspecies (i.e., *C. pusilloides*; *C*. sp. ‘WTT-2016’; Fig. 2). Analysis of the assembled mitogenomes showed subtle nucleotide differences between *C. pusillus* and *C. pusilloides*, such as the coding sequence length for the NAD2 gene, where *C. pusillus* exhibited two amino acid residues longer than *C*. sp. WTT-2013 just prior to the predicted stop codon (Tay et al., 2024). Mating compatibility studies have been applied to demonstrate cryptic species status (Liu et al., 2012; Vyskocilova et al., 2018) and could be consider for *Cryptolestes*.

Using limited microsatellite DNA markers, Toon et al. (2016) reported limited introgression between two highly diverged and geographically separated *C. ferrugineus* partial mtCOI gene lineages, which the authors attributed to historical population structure associated with early Pleistocene climate changes. While introgression between closely related insect species has been shown to be more widespread than previously reported, especially when analysed using genome-wide single nucleotide polymorphic (SNP) markers or via a whole genome resequencing approach (e.g., Anderson et al., 2016, 2018; Elfekih et al., 2021), findings from this study, however, suggested the two *C. ferrugineus* lineages of Toon et al. (2016) likely represented C. *ferrugineus* and *C. pusilloides*, while the likelihood of introgression between closely related *Cryptolestes* species (e.g., *C. pusillus* and *C*. sp. ‘WTT-2013’) remained to be investigated.

The high haplotypes diversity of *C*. sp. ‘WTT-2013’ from Chinese populations and with individuals detected in goods originating from China, suggested that this species likely has an Asian origin. The phosphine resistance status of *C*. sp. ‘WTT-2013’ and of the globally widespread *C*. sp. ‘WTT-2016’ is currently unknown. Phosphine resistance in non-Australian *Cryptolestes* populations has been reported (Daglish, 1999), although the precise species involved will require re-evaluation. Similarly, wide range of phosphine resistance levels in Chinese populations of *C. ferrugineus* have also been reported and may potentially include mixed species (Chen et al., 2023). This work highlights the importance of integrating classical taxonomy with molecular genomic advances, and to survey extensively intra-specific genetic diversity to better gauge nucleotide boundaries for species delimitation. There is a need to revise the taxonomy of *Cryptolestes* and to up-date the existing list of known *Cryptolestes* species to include putative cryptic and novel species (e.g., such as for the potentially misidentified ‘*C. pusillus*’ species (GenBank submitted but unpublished partial mtCOI sequences ON753799.1 -ON753803.1, ON753806.1 -ON753808.1 (accessed date 20-December, 2025) reported in Thailand but which showed the closest match (97.41% sequence identity) to CspWTT2013-Chn10 (Tay et al., 2024)) identified to-date. Future adoption of whole genome HTS methods and associated analysis approaches for *Cryptolestes* species can significantly contribute to agricultural biosecurity and food security preparedness for the stored grain primary industry sector.

## Acknowledgements

WTT and MPC were supported by the Plant Biosecurity Corporative Research Centre project PBCRC2125. DK was supported by CSIRO (R-90035-14). Karl Gordon (CSIRO) and the Department of Agriculture, Fishery and Forestry provided helpful discussion.

